# Extrinsic noise and heavy-tailed laws in gene expression

**DOI:** 10.1101/623371

**Authors:** Lucy Ham, Rowan D. Brackston, Michael P.H. Stumpf

**Author notes:** L.H. and R.D.B. contributed equally to this work.

## Abstract

Noise in gene expression is one of the hallmarks of life at the molecular scale. Here we derive analytical solutions to a set of models describing the molecular mechanisms underlying transcription of DNA into RNA. Our *Ansatz* allows us to incorporate the effects of extrinsic noise – encompassing factors external to the transcription of the individual gene – and discuss the ramifications for heterogeneity in gene product abundance that has been widely observed in single cell data. Crucially, we are able to show that heavy-tailed distributions of RNA copy numbers cannot result from the intrinsic stochasticity in gene expression alone, but must instead reflect extrinsic sources of variability.

---

Trancription is one of the canonical examples of a stochastic process in biology; and as the first step in the central dogma, is of fundamental phenotypic importance. Recent single-cell analysis methods now allow this heterogeneity to be observed in terms of distributions of transcript copy numbers across ensembles of cells. Conveniently, the process is also amenable to the application of elegant mathematical models reminiscent of those in statistical physics Benecke (2008); Schnoerr et al. (2017).

The ***Telegraph process*** is widely used to model stochastic RNA transcription initiation, originally detailed in Peccoud and Ycart (1995) and discussed in recent reviews Munsky et al. (2012); Jones and Elf (2018). In the slightly generalised form we consider here, the gene is either active or inactive: when active, mRNA is transcribed as a Poisson process with rate ***K***_1_, while when inactive, basal transcription may still occur at lower rate, ***K***_0_. The mRNA degradation is modelled by a first-order Poissonian degradation process with rate, *δ*, while switching between the two states occurs at rates *v*_0_ (turn off) and *v*_1_ (turn on), as shown schematically in Fig. 1(a). This leads to a Markov process for the copy number ***n*** of the mRNA molecules at time ***t*** and gene state ***i***, with an associated master equation for the probability ***p***_*i*_(***n***, ***t***),

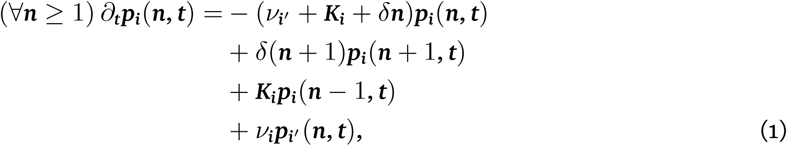

where ***i*** ∈ {0, 1}, and ***i***′ = 1 − ***i***. For, ***n*** = 0 terms involving ***n*** - 1 are set to 0.

**Figure 1:**
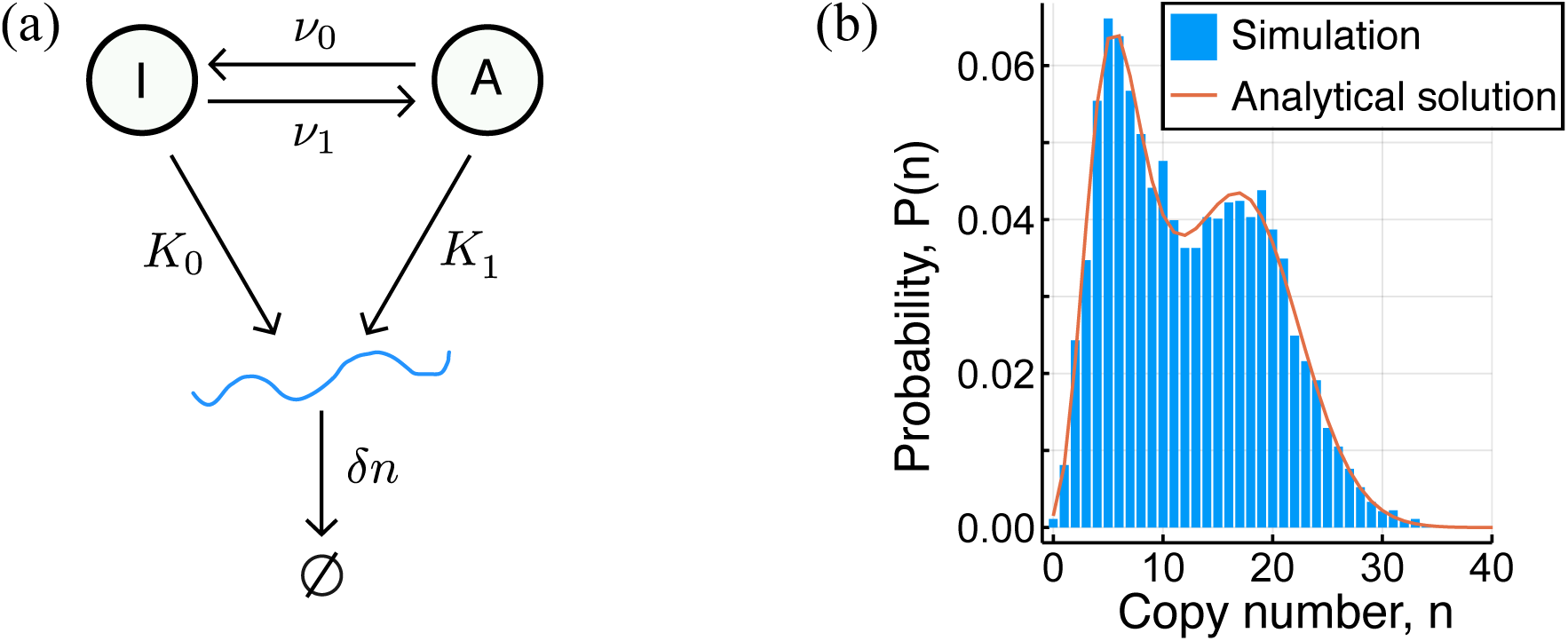
(a) A schematic of the leaky Telegraph model. (b) A comparison of the analytical solution Eq. (3) and a probability mass function from simulated data.

The master equation Eq.(1) for this “leaky gene” model coincides with (Kepler and Elston, 2001, Eq.’s (2),(3)), though a steady state solution is not given there. When ***K***_0_ = 0, the equation reduces to that given in (Peccoud and Ycart, 1995, Eq. (5)). Following the approach of Peccoud and Ycart (1995), generating functions yield an exact solution for the probability generating function (pgf) for the mRNA copy number ^1^:

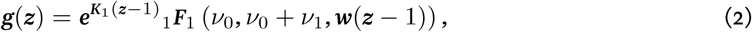

where ***w*** := ***K***_1_ − ***K***_0_ and rates are scaled so that *δ* = 1. The probability mass function (pmf) is then recovered as 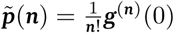, which by the general Leibniz rule gives

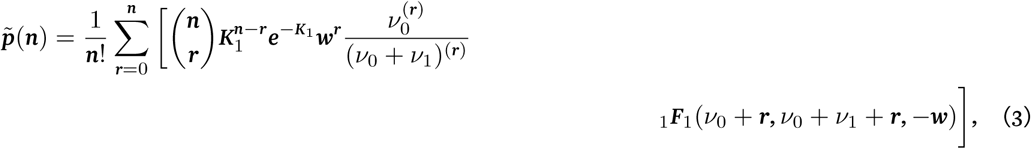

where, for real number ***x*** and positive integer ***n***, the notation ***x***^(***n***)^ abbreviates the rising factorial of ***x***.

A useful limiting case of this generalized model is obtained when the active transcription is sufficiently rare so that transcription can be considered to occur in instantaneous bursts (*v*_0_≫*v*_1_), and the degradation rate is sufficiently small (*v*_0_≫*δ*). This model simultaneously encompasses the two well-known extremes of bursty transcription and constitutive transcription, as we now show.

Under the assumptions *v*_0_≫*δ, v*_1_, the burst size is geometrically distributed, as is already understood Paulsson and Ehrenberg (2000); Ingram et al. (2008):

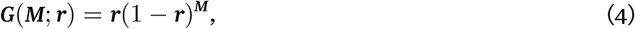

where ***r*** = *v*_0_/(*v*_0_ + ***K***_1_). The master equation Eq. (1) may then be rewritten as,

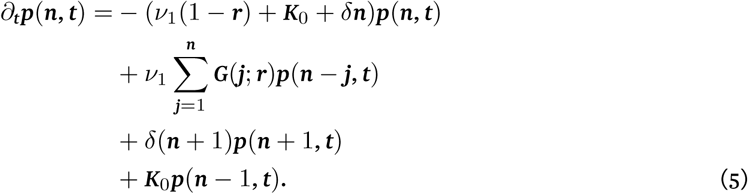

The steady-state solution is again obtained using generating functions, with the exact expression for the pgf given by

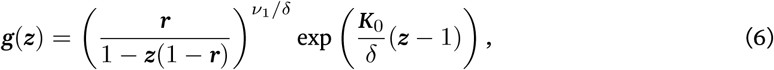

which is the product of the pgf for the negative binomial distribution NegBin(*v*_1_/*δ*, ***r***) and the Poisson distribution Pois(***K***_0_/*δ*). Thus **?**,

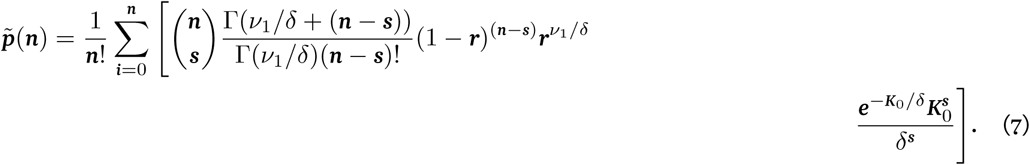

When ***K***_0_ = 0, corresponding to bursty gene expression, Eq. (S.23) is the pgf for NegBin(*v*_1_/*δ*, ***r***), agreeing with the solution of Paulsson and Ehrenberg (2000). Alternatively, use Eq. (7), using 0^0^ := 1. When ***K***_1_ = 0 (or indeed is kept constant and *v*_0_ → *∞*) the burst height parameter ***r*** = *v*_0_/(*v*_0_ + ***K***_1_) becomes 1, and the steady state solution Eq. (7) agrees with the solution for constitutive gene expression, namely the Pois(***K***_0_/*δ*) distribution; see Fig. 2 for the pdfs of both of these.

**Figure 2:**
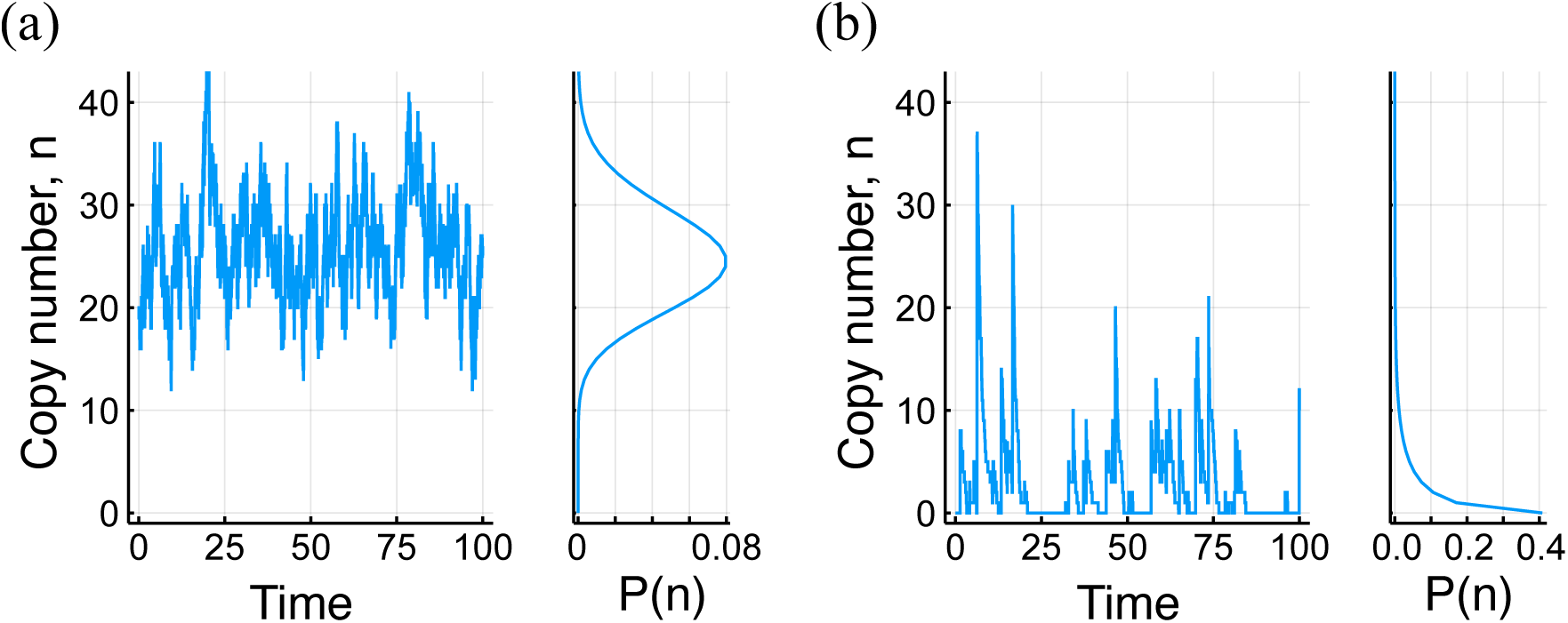
Time series and corresponding probability distributions for (a) constitutive and (b) bursty transcription.

Similarly, when ***K***_0_ = 0, the generating function Eq. (S.13) for the leaky gene model reduces to the solution for ***g***(***z***) in (Peccoud and Ycart, 1995, Eq. 20) by way of Kummer’s transformation (Olver et al., 2010,§13.2.39). This then yields the following well-known analytical expression for the Telegraph model Peccoud and Ycart (1995); Raj et al. (2006)

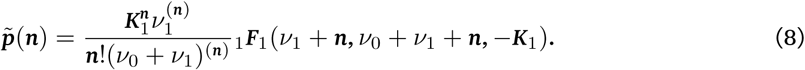

The standard Telegraph model for transcription describes the effect of intrinsic noise at the level of a single gene, yet the process will often also be influenced by other sources of variability. Such extrinsic noise has been widely observed experimentally Elowitz (2002); Swain et al. (2002); Raser and O’Shea (2004); Gasch et al. (2017), and considered theoretically Dattani and Barahona (2016); Fu and Pachter (2016); Bressloff and Levien (2019); Thomas (2019), but incorporating these effects into the Master equations has generally proven challenging. The approach we take in the context of the Telegraph model is to consider the model parameters themselves to vary between cells, and therefore to be drawn from probability distributions Lenive et al. (2016). The mRNA copy number then follows a compound distribution,

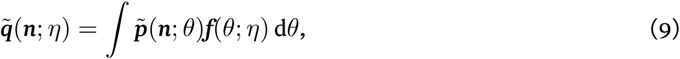

where *θ* is the vector of parameters [*v*_0_, *v*_1_, ***K***_0_, ***K***_1_] and the distribution ***f*** is a multivariate distribution for *θ* with hyperparameters *η*. This model is valid provided that parameter values are static for individual cells but vary across an ensemble of cells according to ***f***, or if the parameter values are dynamic, but change at substantially slower timescales (adiabatically) relative to the transcriptional dynamics. An example of the latter is variation in upstream biological drivers, which is a special case of the extrinsic noise considered in Dattani and Barahona (2016). For the remainder of the paper, when it is clear from the context that only one rate of transcription is being considered, we will use ***K*** in place of ***K***_0_ or ***K***_1_.

An interesting example arises from Eq. (8) when ***K*** is Gamma(*α, β*) distributed. In this case, the compound distribution 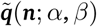 can be shown to be

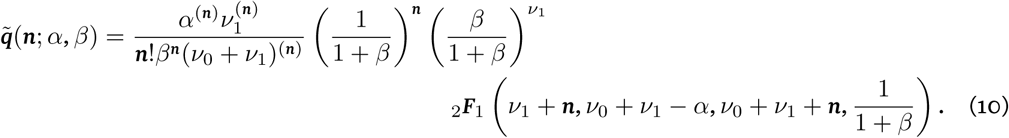

Rather intriguingly, this distribution is also the steady-state protein number distribution found in Shahrezaei and Swain (2008). Gene expression there is modelled as a three-stage stochastic process, but the model can be considered as Telegraph noise, with distribution Eq. (8), on the rate parameter of a Poissonian process. It is surprising that the resultant steady-state distribution coincides with that of Gamma-distributed noise on a Telegraph process.

Another striking example arises by way of constitutive transcription 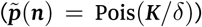 again with *K ∼* Gamma(*α, β*). In this case, the compound distribution yields the negative binomial distribution,

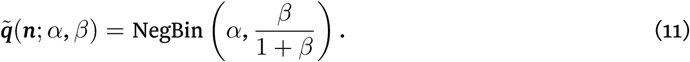

Thus the copy number distribution alone cannot distinguish constitutive expression with extrinsic noise, from a bursty expression without extrinsic noise, which has distribution NegBin (*v*_1_/*δ*, ***r***). A similar example is given in Dattani and Barahona (2016), where constitutive expression with Beta-distributed ***K*** is shown to agree with the solution of the Telegraph model, Eq. (8).

Whether or not the moments of compound distributions converge is a problem of considerable practical importance Willinger et al. (2004). We can provide simple formulæ for all moments of the copy number distribution Eq. (8) under extrinsic noise on the transcription rate ***K***. Noise in ***K*** is of particular relevance, as will become apparent in due course. Let ***X*** = ***X*_*K*_** denote a random variable from 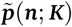 (from Eq. (8)) and ***Y*** = ***Y***_*η*_ a random variable from the compound distribution 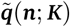 (from Eq. (9)). It can be shown that the ***n***^th^ moment of ***Y*** is given by

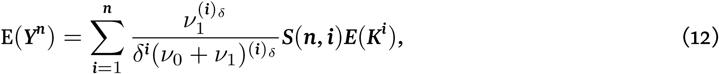

where ***S***(***n***, ***i***) is a Stirling number of the second kind; the notation 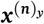 abbreviates ***x***(***x***+***y***) … (***x***+***y***(***n*** - 1)) for real numbers ***x***, ***y*** and positive integer, ***n***. It follows from Eq. (S.31) that if the first two moments of the compounding distribution ***f***(***K***; *η*) are known, then the mean, variance and Fano factor of the compound distribution 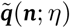 can be easily calculated. From Eq. (S.31), and noting that ***S***(1, 1) = ***S***(2, 1) = ***S***(2, 2) = 1, the Fano factor of 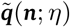 is given by

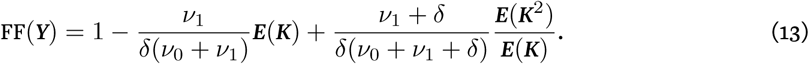

It is also possible to obtain formulæ for the ***n***^th^ moments and Fano factor in the case for constitutive expression with extrinsic noise on ***K***, as well as in the case for bursty expression with extrinsic noise on ***K***. In the former, we can alternatively obtain a formula for the Fano factor from Eq. (13) by taking 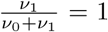 (corresponding to *ν*_1_ → *∞*, or *ν*_0_ → 0). This gives

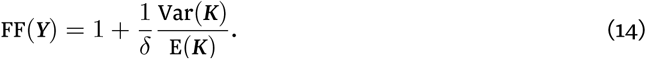

It has previously been suggested that heterogeneity in mRNA copy numbers and a universal scaling between the mean and Fano factor is attributable primarily to intrinsic noise, as opposed to extrinsic noise So et al. (2011). The purported observational evidence for these claims is: (i) in the limit of low mean mRNA copy number, the Fano factor is approximately one; (ii) at the other extreme of high mean mRNA copy number, the Fano factor decreases sharply rather than approaching a plateau; (iii) the square of the coefficient of variation decreases monotonically with the mean.

These claims do not stand up against the analytical solutions for the Fano factor obtained above. From Eq. (13), the Fano factor of the compound distribution 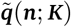 depends only on E(***K***) and Var(***K***) (or equivalently E(***K***^2^)) and the values of the parameters *ν*_0_, *ν*_1_, *δ*. Throughout we let *c* denote the coefficient of variation, *σ*(***K***)/ E(***K***), for the noise distribution, ***f***(***K***; *η*). Noting that E(***K***^2^)/ E(***K***) = (1 + ***c***^2^) E(***K***), Eq. (13) becomes

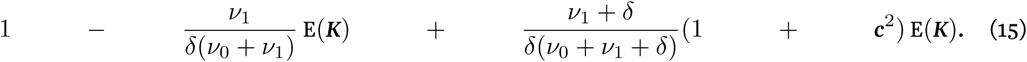

The situation ***E***(***Y***) → 0 corresponds to *ν*_0_ → *∞*, which from Eq. (15) easily gives FF(***Y***) → 1. Thus, item (i) holds true for all values of ***c***, and therefore for any extrinsic noise distribution on K, provided that the second moment exists.

Figure 3(a) shows qualitatively identical behaviour of the Fano factor as a function of the mean mRNA copy number for values of *c* close to 1, even for the same parameter values of *ν*_1_, *δ*, ***E***(***K***) = ***K***. We remark that, as in So et al. (2011), the mean is varied by regulating *ν*_0_ only. Values of *c* below 0.6 have been omitted, as they are visually indistinguishable from the intrinsic noise case at the present resolution.

**Figure 3:**
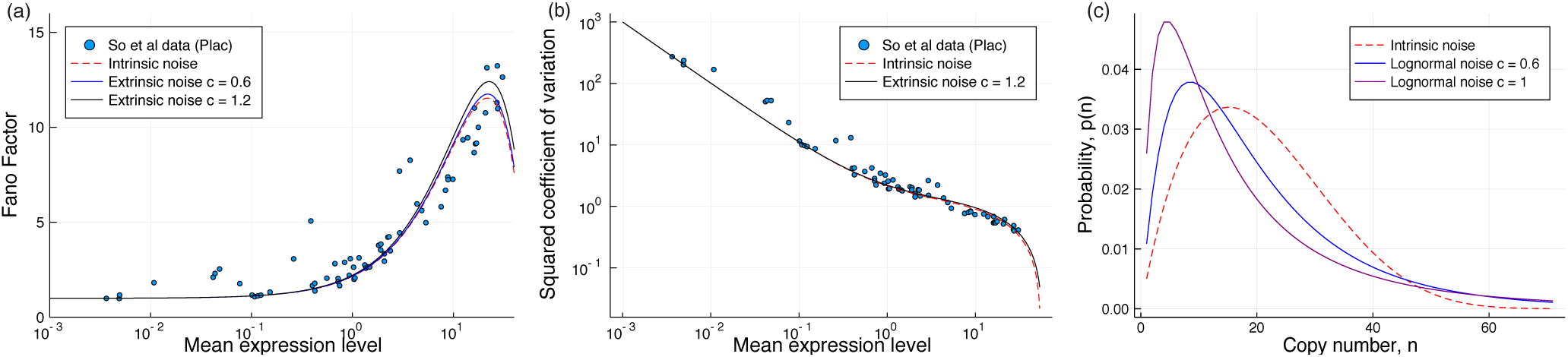
In figures (a) and (b), the parameters are E(K) = K = 54, v_1_ = 0.83 and δ = 1. (a) The Fano factor of the compound distribution as a function of the mean mRNA copy number (given by Eq. (15)), varied by tuning the parameter v_0_. Different values of c are plotted against the universal noise scaling curve (intrinsic noise only) given in So et al. (2011). Data is the P_lac_ promoter also from So et al. (2011). (b) The squared coefficient of variation as a function of the mean copy number for intrinsic and extrinsic noise c = 1.2. (c) The effect of lognormal extrinsic noise on K on the copy number distribution, with K = 60, v_0_ = 4, v_1_ = 2 and δ = 1.

Finally, we consider the squared coefficient of variation as a function of the mean copy number for different values of ***c***; see Fig. 3(b), and note that values of ***c*** smaller than ***c*** = 1.2 are again visually (and practically) indistinguishable from intrinsic noise and so are omitted. We see that the behaviour is effectively identical to that of intrinsic noise only. Even for unrealistic values of ***c*** (for example ***c*** = 200, 000), the squared noise curve continues to satisfy (iii).

We next examine the effect of extrinsic noise on the noise scaling curve in the case for constitutive transcription, again considering only noise on ***K***. In this case, it can be easily shown that the mean copy number E(***Y***) is equal to E(***K***). Thus, from Eq. (14), scaling the Fano factor with mean copy number is dependent only on the noise distribution of ***K***. If the coefficient of variation is fixed at ***c*** as the noise distribution on *K* is varied, then the Fano factor is given by 1 + ***c***^2^ E(***K***), which is linear in E(***Y***) = E(***K***). On semilogarithmic axes and plotted as a function of the mean copy number, this yields the same qualitative behaviour as that found in (Jones et al., 2014, Fig. 2B). The observations after Eq. (11) are pertinent here: with 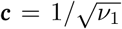, identical noise scaling behaviour arises from the two extremes—constitutive expression with extrinsic noise and bursty expression without noise.

Thus far, extrinsic noise, as modelled by the compound distribution, exhibits behaviours that are similar to intrinsic noise alone. We now present a potential qualitative identifier for extrinsic noise: we show that contrary to previous claims Iyer-Biswas et al. (2009), intrinsic noise alone never leads to a heavy-tailed copy number distribution, but find many cases in which extrinsic noise does so. Formally, we take heavy tailed to mean that the moment generating function (mgf) is undefined for positive ***t***, which implies that the tail of the distribution decays more slowly than that of the exponential distribution.

If *K* and *δ* are fixed, then the copy number is maximized when the gene remains permanently active, which has distribution Poisson(***K***/*δ*) and is not heavy tailed. Intuitively then, no compounding of *ν*_0_ and *ν*_1_ alone results in a heavy tailed distribution. A more robust argument is obtained by establishing the following inequality for the Telegraph model **?:** for all positive ***t***,

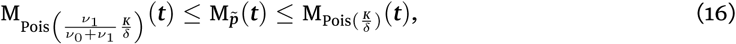

where M_*g*_ denotes the mgf for distribution *g*. In particular, 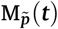 is bounded above by a Poissonian mgf that does not depend on *ν*_0_ or *ν*_1_. Thus 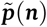 itself is not heavy tailed, and we require compounding of ***K*** or *δ* to make it so. On the other hand, any extrinsic noise on ***K*** or *δ* that renders the mgf for Pois 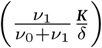 undefined, will also result in 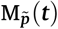 being undefined and the resulting compound distribution will be heavy tailed. We now demonstrate this for ***K*** *∼* LogNormal(*µ, σ*). The result relies on the following well-known property of mixture distributions, here interpreted in the context of Eq. (9):

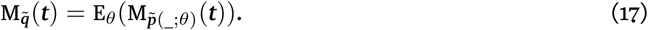

It follows from this and Eq. (16) that the compounding integrand is bounded below by

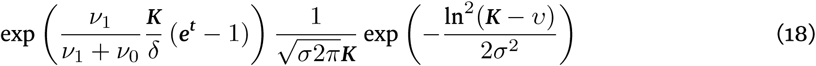

which diverges to infinity as ***K*** → *∞* provided ***t*** *>* 0. Thus log-normal extrinsic noise on ***K*** renders the compound distribution 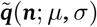 heavy tailed; cf. Fig. 3 (c). These results extend to the leaky gene model, with log-normal noise on ***K***_1_ (conditional on ***K***_0_ *<* ***K***_1_), as it is trivial that ***K***_0_ *>* 0 only increases the probability of large copy number in comparison to the standard Telegraph model.

From Eq. (16) we see that when 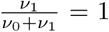 (that is, *ν*_1_ → *∞*, or *ν*_0_ → 0) we obtain the constitutive distribution, Poisson(***K***/*δ*). When ***K*** is subject to log-normal noise, the argument using Eq. (18) carries through unchanged.

For bursty expression we require *ν*_0_ ≫ *ν*_1_, *δ* so consider extrinsic noise on ***K*** only. The effect of extrinsic noise here is qualitatively different to the other cases: we observe that the mgf for the negative binomial distribution is given by,

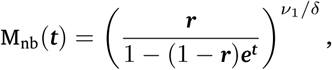

for ***t*** *< -*ln(1*-****r***) = *-*ln(***K***/(***K*** + *ν*_0_)), and is infinite otherwise, where ***r*** = *ν*_0_/(*ν*_0_ + ***K***). Thus, the range of positive ***t*** for which M_nb_(***t***) is finite approaches 0 as ***K*** → *∞,* implying that any unbounded distribution ***f***(***K***; *η*) results in the moment generating function of the compound distribution

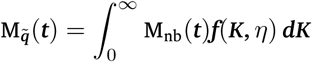

undefined for positive ***t***. Now in contrast to the previous result, Eq. (11), Gamma distributed noise on ***K*** yields a heavy-tailed distribution.

In summary, our approach has provided a range of new analytical solutions to long-standing problems in gene expression modelling. Crucially, we have been able to include extrinsic noise and found that this alone provides complementary and alternative explanations for many empirical observations So et al. (2011). Further to this, we have demonstrated that extrinsic noise can explain observations of heavy-tailed distributions Bengtsson (2005), which intrinsic noise alone cannot. Given the notoriously noisy environment within cells and the intricate organisation of gene regulatory networks, noise extrinsic to a given gene (or gene model) is almost certainly ubiquitous. The framework and results provided here allow us to get a better, more detailed handle on the origins and implications of noise in molecular systems and beyond.

## Acknowledgments

LH and MPHS gratefully acknowledge funding from the University of Melbourne DVCR fund; RDB and MPHS are funded from the BBSRC through Grant No. BB/N003608/1.

L.H. and R.D.B. contributed equally to this work.

## A. The leaky gene model

### A.1 Steady-state solution

In this section, we provide a full derivation of the steady-state solution to the leaky gene model described by the following master equation, given as Eq. 1 in the main text.

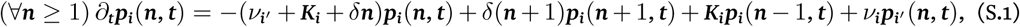

where *i* ∈ {0, 1},, and *i′* = 1–*i*. For *n* = 0, terms involving *n–*1 are set to 0. As mentioned, the development closely follows Peccoud and Ycart (1995). Letting *p*_*i*_(*n*) denote the steady-state probability of copy number *n* in state *i* ∈ {0, 1}, we introduce the following generating functions:

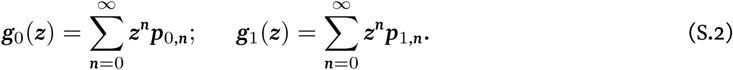

Multiplying Equation (S.1) through by *z*^*n*^ and then summing these over *n* from 0 to *∞* gives rise to the following pair of symmetric first-order differential equations:

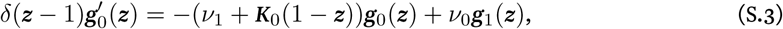

and

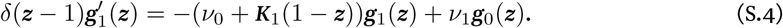

By differentiation, then back substitution, we arrive at the following symmetric pair of second-order differential equations:

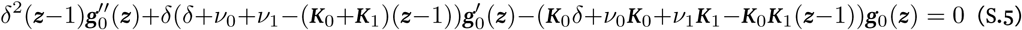

and

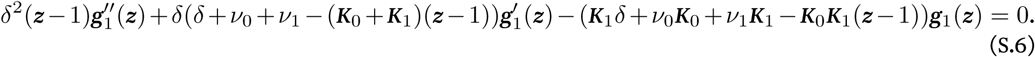

As in Peccoud and Ycart (1995), the substitution *x* = 1 – *z* and defining *h*_*i*_(*x*) = *g*_*i*_(1 – *x*), for *i* ∈ {0, 1}, leads to conceptually simpler equations

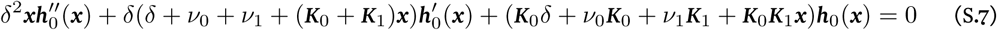

and

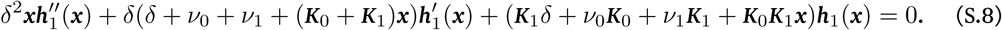

In comparison to Peccoud and Ycart (1995), these equations have an extra term in the *h*_*i*_(*z*) coefficient which makes the solution more complicated. Nevertheless, both (S.7) and (S.8) can be represented as extended confluent hypergeometric equations and so have a known general solution. At this point it is notationally convenient to set *δ* = 1, which is equivalent to scaling time by *δ* and there is no loss of generality in doing this. Let

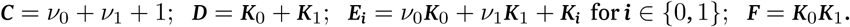

Then from (S.7) and (S.8), we obtain the equations

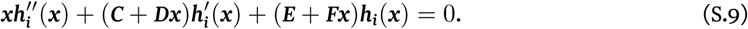

Given the initial conditions *h*_0_(0) = *v*_0_/(*v*_0_ + *v*_1_) and *h*_1_(0) = *v*_1_/(*v*_0_ + *v*_1_), these can be solved as

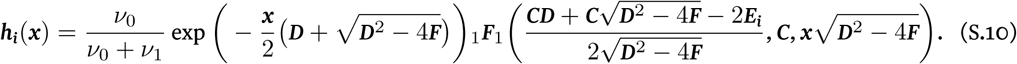

Note that (S.7) and (S.8) have a second solution in terms of Kummer’s second hypergeometric function, however this is not defined at the initial conditions and fails other physical properties of the system; this is detailed also in Grima et al. (2012). We observe that this solution uses the assumption *K*_0_ ≠ *K*_1_, however the special case of *K*_0_ = *K*_1_ is a degenerate form of the system (a standard birth-death model). Thus we assume now that *K*_0_ *< K*_1_, while the o<u>pposite</u> case *K*_1_ *< K*_0_ follows by symm<u>etry. Th</u>is assumption leads to enormous simplification, because *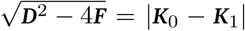* and then 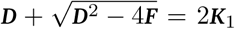. After further cancelling in (S.10), and replacing *x* by 1 *- z* we obtain

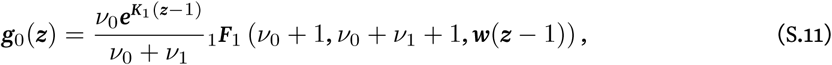

and

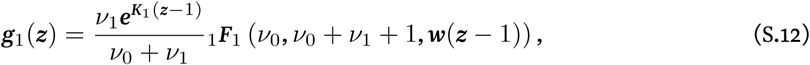

where *w* denotes *K*_0_*-K*_1_. For the copy number itself, we want *g*(*z*) = *g*_0_(*z*) + *g*_1_(*z*). Using the functional identity (Olver et al., 2010, §13.3.3) for the confluent hypergeometric function

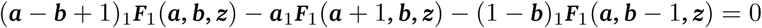

we obtain

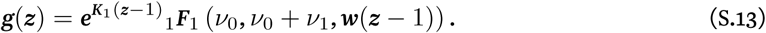

The probability mass function is then recovered by way of 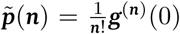. The general Leibniz rule now gives

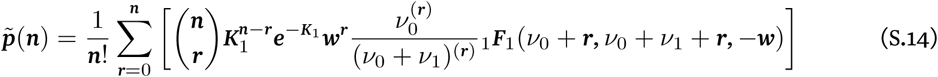

### A.2 A special case

We now provide a full derivation of the solution to the limiting case of the leaky gene model; refer to the master equation Eq. 5 in the main article, or alternatively see (S.20) below. Recall that this model is justified only when the active transcription is rare enough to be considered instantaneous (*v*_0_ ≫*v*_1_) and the degradation rate is sufficiently small (*v*_0_ ≫*δ*). Consider then an interval of length *T* in which the gene is active at the higher rate *K*_1_ and in which no degradation occurs. The probability of transcribing *M* mRNA molecules is given by a Poisson distribution as,

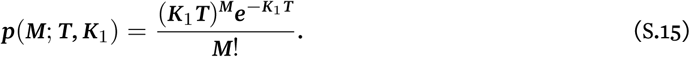

The time period *T* is itself exponentially distributed according to,

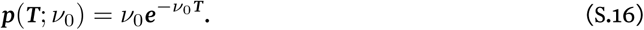

Marginalising (S.15) over *T* leads to,

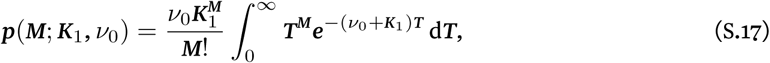

which with the substitution *u* = (*v*_0_ + *K*_1_)*T* can be solved to yield,

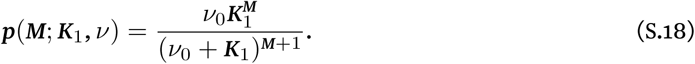

As is already understood, this is the geometric distribution Paulsson and Ehrenberg (2000); Ingram et al. (2008),

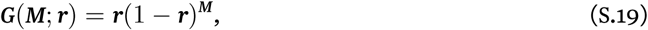

where *r* = *v*_0_/(*v*_0_ + *K*_1_). Given instantaneous geometric bursts of transcription, the master equation (S.1) may then be rewritten as,

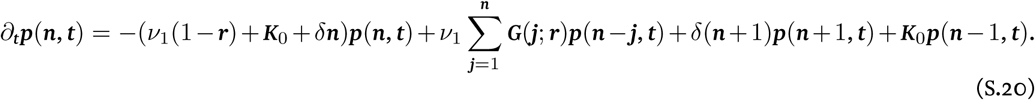

The first term in this equation gives fluxes away from *p*(*n*), occurring due to bursts of non-zero size or degradation of a single mRNA. The second term gives the total probability of aburst occurring at any lower molecule number and raising the number to *n*, while the third term gives the probability of degradation reducing the molecule number from *n* + 1. The fourth term is the probability of raising the number to *n* when inactive. To solve this system in the steady state we introduce the following generating function:

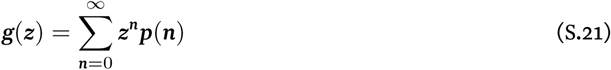

Multiplying (S.20) through by *z*_*n*_ and summing over *n* from 0 to *∞* gives the following equation:

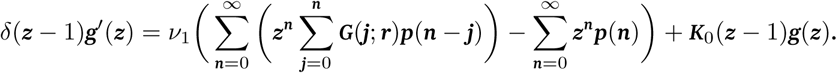

which after regrouping becomes

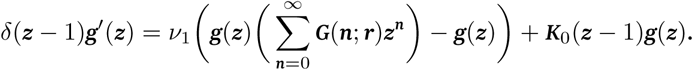

The term 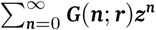 is the probability generating function for the geometric distribution, which can be written as 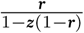. Thus we obtain the first-order differential equation

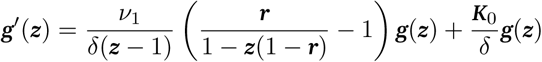

which simplifies to

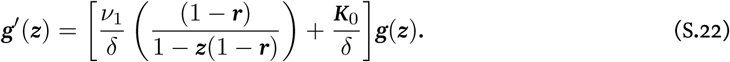

As previously mentioned, this is routinely solved by the integrating factor method (using the initial condition *g*(1) = 1) to give

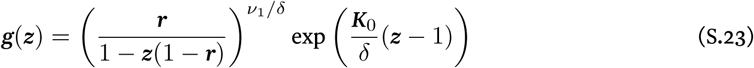

Which is the product of the probability generating function for the negative binomial distribution NegBin(*v*_1_/*δ, r*) and the Poisson distribution Pois(*K*_0_/*δ*). This gives the following analytical expression for the steady-state solution, presented as Eq. 7 in the main article:

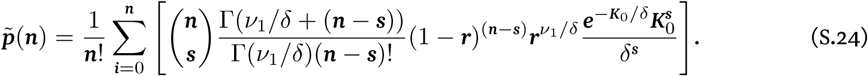

## B Incorporating extrinsic noise

As mentioned in the main article, an interesting compound distribution arises from Eq. 8 when *K* is distributed according to the distribution Gamma(*α, β*); see Eq. 10. We now provide the missing details of its derivation. From Eq. 9, the compound distribution 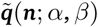 is given by

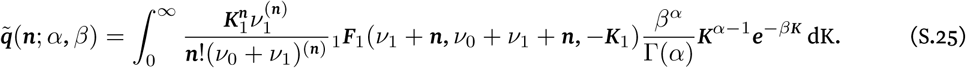

Using the Laplace transformation (Olver et al., 2010, §13.10.3),

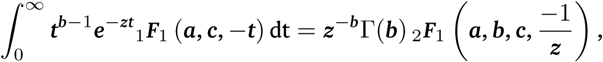

the Eq. (S.25) becomes

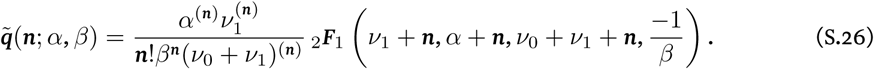

Now applying the linear transformation (Olver et al., 2010, §15.8.1)

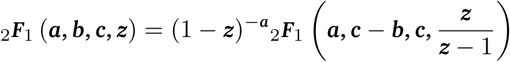

to (S.26), we obtain

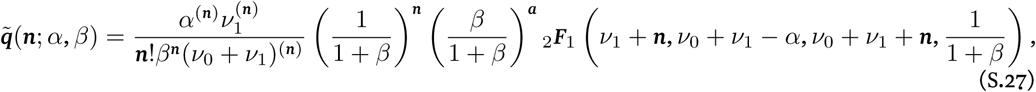

## C Existence of moments under extrinsic noise

In the main article we provide simple formulæ for the ***n***^th^ moments of copy number distributions under extrinsic noise on the rate parameter ***K***. The following three subsections are dedicated to obtaining these: the first covers the Telegraph model under extrinsic noise, and the second and third cover constitutive and bursty expression under extrinsic noise, respectively. Throughout, we let ***f***(***K***; *η*) denote the compounding distribution of *K*.

### C.1 Telegraph model

Let ***X* = *X***_***K***_ denote a random variable with distribution 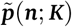 (from Eq. 8) and ***Y* = *Y***_*η*_ a random variable with the compound distribution 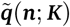 (from Eq. 9). In the following it will be useful to recall that, for real numbers ***x***, ***y*** and positive integer ***n***, the notation 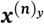 abbreviates ***x***(***x*** + ***y***) … (***x*** + ***y***(***n*** − 1)), the rising factorial of ***x*** with respect to *δ*. The notation ***x***_(***n***)_ abbreviates ***x***(***x*** − 1) … (***x*** − (***n*** − 1)), the falling factorial of ***x***. Our results rely on the following property of mixture distributions.

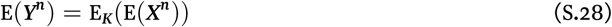

The proof of (S.28) is straightforward, relying only on switching the order of integration/sum. Following Peccoud and Ycart Peccoud and Ycart (1995), we let

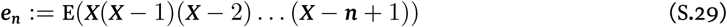

and obtain directly from the generating function (Peccoud and Ycart, 1995, Equation 20),

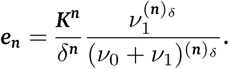

Before we continue, we recall that Stirling numbers of the first and second kind relate expansions of the falling factorial ***x***_(***n***)_ := ***x***(***x*** − 1) … (***x*** − ***n*** + 1) to conventional powers of ***x***. Stirling numbers of the first kind are the coefficients in the expansion of ***x***_(***n***)_, and Stirling numbers of the second kind arise as the coefficients when ***x***^***n***^ is written as a linear combination of ***x***_(1)_, …, ***x***_(***n***)_. Thus, from (S.29) we have that 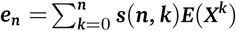, where ***s***(***n***, ***k***) are Stirling numbers of the first kind. We obtain

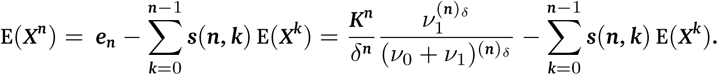

Thus E(*X*^***n***^), as a function of ***K***, is a polynomial of degree ***n***, which we can write as

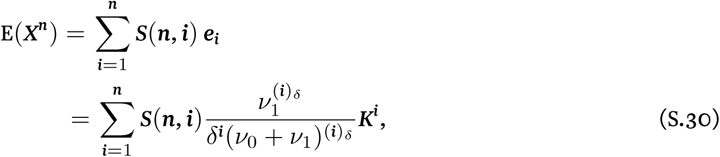

where ***S***(***n***, ***i***) is a Stirling number of the second kind. Returning to (S.28), we obtain

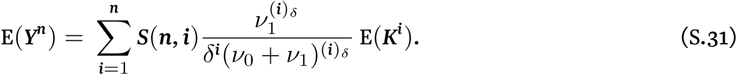

Noting that ***S***(1, 1) = ***S***(2, 1) = ***S***(2, 2) = 1, it follows from (S.31) that

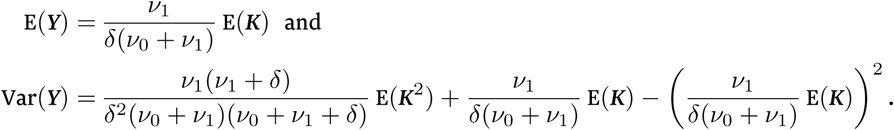

This then yields the following formula for the Fano factor of 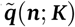:

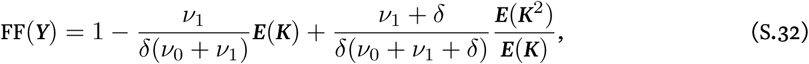

which is presented as Eq. 13 in the main article.

### C.2 Constitutive expression

In a similar way to the Telegraph model, we can obtain closed-form formulæ for the ***n***^th^ moments and Fano factor in the case for constitutive expression with extrinsic noise on ***K***. Let ***X* = *X***_***k***_ be a random variable with distribution Pois(***K***/*δ*) distribution, let 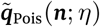 denote the compound distribution obtained by compounding Pois(***K***/*δ*) by ***f***(***K***; *η*), and let ***Y* = *Y***_*η*_ denote a random variable for 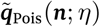. The ***n***^th^ moments of a Pois(***K***/*δ*) random variable are given by

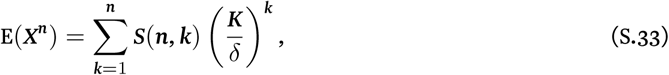

where ***S***(***n***, ***k***) is a Stirling number of the second kind. From (S.28), it follows immediately that

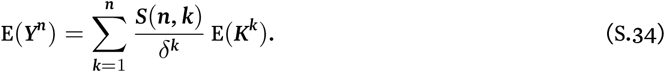

Thus,

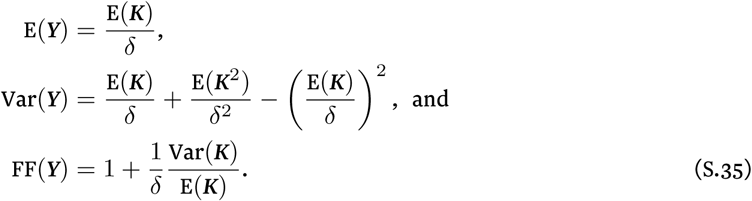

### C.3 Bursty expression

Let ***X* = *X***_***K***_ denote a random variable with distribution the NegBin(*ν*_1_/*δ*, ***r***) distribution, where ***r*** is equal to *ν*_0_/(*ν*_0_ + ***K***). Let 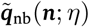 denote the compound distribution obtained by compounding NegBin(*ν*_1_/*δ*, ***r***) by ***f***(***K***; *η*), and let *Y* = *Y*_*η*_ denote a random variable for 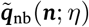. Recall that the moment generating function for the NegBin(*ν*_1_/*δ*, ***r***) distribution is given by

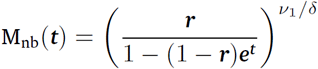

for ***t*** *<* − ln(1 − ***r***) = − ln(***K***/(***K*** + *ν*_0_)), and is infinite otherwise. Using the recurrence relation,

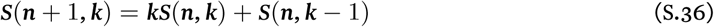

a straightforward proof by induction on the number ***n*** of derivatives of M_nb_(***t***) with respect to ***t*** (with base case ***n*** = 0 given by M_nb_(***t***)) shows that

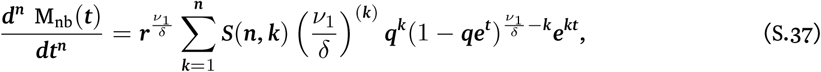

where ***q*** = 1 − ***r***. By substituting ***t*** = 0 into (S.37), we immediately obtain

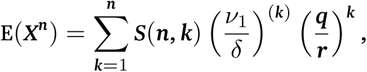

which simplifies to

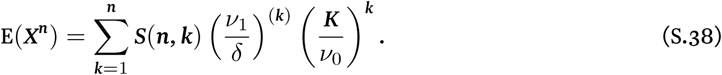

It then follows from (S.28) that

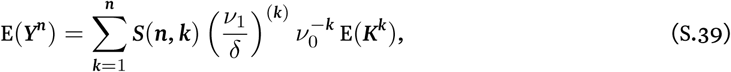

which gives the following formula for the Fano factor of 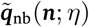:

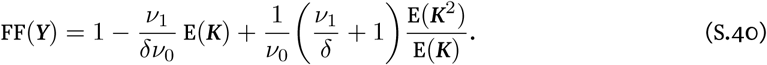

## D Heavy-tailedness of the compound distribution

In the final section of the main article we showed that heavy-tailedness in the copy number distribution is a potential qualitative identifier of extrinsic noise. The arguments rely on the following inequality for the Telegraph model, given as Eq. 16 in the main article: for all positive ***t***,

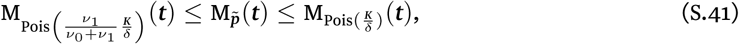

where M_***g***_ denotes the mgf for distribution ***g***. We now give a detailed proof of this result. In the following, let 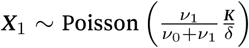, let ***X***_2_ ∼ Poisson 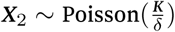, and let 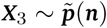 (from Eq. 8). For ***n*** ∈ℕ, we have

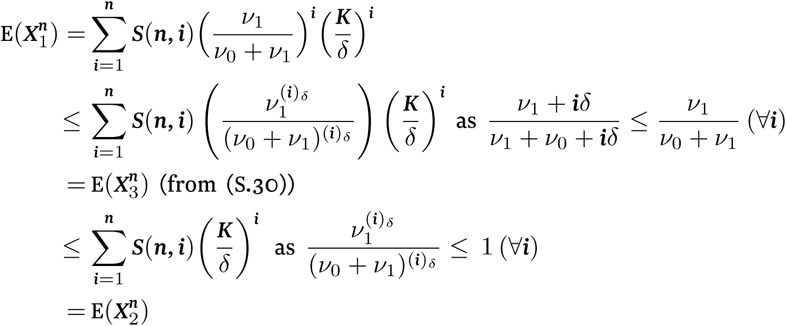

As each moment 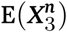 is sandwiched between two Poissonian ***n***^th^ moments, it immediately follows from the least upper bound property that the mgf for the copy number distribution 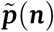 is bounded below by the mgf for the Pois 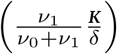 distribution and bounded above by the mgf for the Pois 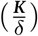 distribution. Thus, Equation (S.41) holds.

## Footnotes

1 See Supplementary Material at [URL] for derivations of the steady-state solutions to the leaky gene model, Eq. 3, and its limiting case, Eq. 7, the derivation of the Fano factor under extrinsic noise, Eq. 13, and a proof of the inequality, Eq. 16, used in the arguments for heavy tailed-ness of copy number distributions under extrinsic noise. Includes Refs. Peccoud and Ycart (1995); Grima et al. (2012); Olver et al. (2010); Paulsson and Ehrenberg (2000); Ingram et al. (2008).

